# The MOS4-associated complex subunit MAC5A maintains meristem development by regulating transcription elongation in Arabidopsis

**DOI:** 10.64898/2026.08.27.747678

**Authors:** Meng Ye, Xudong Li, Yufeng Zhou, Xiaojuan Huang, Huilin Liu, Shuqing Liu, Ruibo Hu, Aixia Li, Shengjun Li

**Affiliations:** Qingdao Institute of Bioenergy and Bioprocess Technology, Chinese Academy of Sciences, Qingdao 266101, China; University of Chinese Academy of Sciences, Beijing 100049, China; State Key Laboratory of Wheat Improvement, College of Agronomy, Shandong Agricultural University, Tai’an 271018, China; College of Agriculture and Bioengineering (Peony College), Heze University, Heze 274015, China

**Author notes:** Corresponding author: Shengjun Li College of Agronomy, Shandong Agricultural University, Tai’an, China.

**Keywords:** Transcriptional elongation, pre-mRNA splicing, meristem maintenance, MAC5A, ELP6

## Abstract

The precise coordination of co-transcriptional RNA processing with transcriptional elongation is essential for eukaryotic gene regulation, yet the machineries coupling these processes remain largely elusive. Here, we reveal a role for the spliceosome-associated MOS4-associated complex (MAC) in regulating plant development through the direct regulation of transcriptional elongation. We demonstrate that the MAC subunit MAC5A physically interacts with Elongator Protein 6 (ELP6), a core component of the transcriptional Elongator complex. Genetic analyses reveal that MAC5A and ELP6 function synergistically to regulate apical meristem activity. Importantly, the MAC5A-Elongator module is required for the efficient expression of the critical auxin efflux carrier PIN-FORMED1 (PIN1) by enhancing RNA Polymerase II (RNAPII) occupancy across the *PIN1* locus. Furthermore, we show that this elongation-promoting function is not unique to MAC5A, as other core MAC components are similarly required for efficient transcription progression. Together, our findings uncover a splicing-elongation nexus where MAC5A likely acts as a molecular bridge between the spliceosome and the elongation polymerase. This functional coupling ensures the efficient transcription of key developmental regulators, providing a mechanistic framework for the coupling of RNA processing to transcription and suggesting a conserved principle of gene expression control across eukaryotes.

**Significance:** The RNA-binding protein MAC5A plays an essential role in RNA metabolism in eukaryotes. This study shows that MAC5A functions as a molecular adaptor connecting the MOS4-associated complex to the Elongator complex, thereby coordinating efficient transcription of the auxin transporter *PIN1* to sustain auxin flow and meristem maintenance in Arabidopsis. This discovery establishes a new conceptual framework for co-transcriptional regulation in plant development and provides broad insights into the molecular control of growth across plant species.

## Introduction

The phytohormone auxin is a master regulator of plant development, governing fundamental processes from organ initiation and spatial patterning to the maintenance of stem cell niches within meristems (1–3). Developmental coordination relies on the establishment of precise local auxin maxima and gradients within tissues, primarily achieved through the polar localization of PIN-FORMED (PIN) efflux carriers (4–7). Plants maintain these auxin distributions via a tightly regulated network encompassing auxin biosynthesis, transport, and signaling (8) (9–11).

This regulatory network operates at multiple levels. At the transcriptional stage, transcription factors play a well-established role in initiating the expression of auxin-related genes. For instance, MONOPTEROS (MP) directly binds to the promoters of *PIN1*, *PIN3*, and *PIN7* to activate their transcription (12), while the MADS transcription factor XAANTAL2 (XAL2) specifically binds to the promoters of *PIN1* and *PIN4* to coordinate auxin transport (13). Chromatin remodelers further refine transcriptional activation. For example, MP recruits the SWI/SNF chromatin remodeling ATPase BRAHMA (BRM) and SPLAYED (SYD) to enhance DNA accessibility at target loci (14, 15). While transcriptional initiation is increasing well-characterized, the subsequent step of transcription elongation and its contribution to auxin-mediated development remain poorly understood.

In eukaryotes, RNA polymerase II (RNAPII)-mediated transcriptional elongation is a highly dynamic and tightly regulated process essential for precise gene expression (16–18). The evolutionarily conserved Elongator complex facilitates elongation by associating with elongating RNAPII. Elongator comprises a catalytic core (ELP1-ELP3) and an accessory subcomplex (ELP4-ELP6) critical for substrate recognition and structural stability (19, 20). In plants, Elongator contributes to development by affecting RNAPII-mediated transcription elongation of auxin-related genes (20), with mutants exhibiting severe defects, including short roots and narrow leaves (21–23). Elongator also controls tRNA modification, influencing auxin distribution and responsiveness, suggesting its multiple roles in plant growth and development (24). Nevertheless, the precise molecular mechanisms by which Elongator regulates transcriptional rates, particularly its recruitment and interaction with transcriptional cofactors, remain largely enigmatic.

In eukaryotic systems, gene transcription is tightly coupled to RNA processing, including pre-mRNA splicing, in a co-transcriptional manner. This coupling ensures the coordinated and efficient gene expression. The MOS4-associated complex (MAC) is an evolutionarily conserved multi-protein complex, functioning in pre-mRNA splicing and miRNA biogenesis, with key roles in development and stress responses (25, 26). The accessory subunit MAC5 has been demonstrated to be crucial for DNA damage repair, miRNA biogenesis, and disease resistance in Arabidopsis (26–28). The genetic redundancy between its two encoding genes, *MAC5A* and *MAC5B*, is unequal. Between them, MAC5A serves as the dominant contributor, as evidenced by the severe developmental defects observed in the *mac5a MAC5B*/*mac5b* mutant, including narrow, pin-like leaves and complete sterility (26, 28).

Despite MAC5’s established importance in RNA-related processes, its potential role in hormonal regulation, particularly in auxin-mediated development, has remained unexplored. In this study, we unveil a novel function of MAC5A in auxin-dependent plant development. We demonstrate that MAC5A physically interacts with ELP6, an accessory subunit of the Elongator complex. Furthermore, we establish that MAC5A and ELP6 act synergistically to regulate apical meristem development by modifying the transcriptional elongation of the auxin efflux carrier gene *PIN1*. Collectively, our findings not only assign a new developmental role to MAC5A but also provide a mechanistic link between a key splicing-associated complex and the Elongator in the transcriptional control of a central auxin transporter, suggesting its significance in the adaptive evolution of plant architecture.

## Results

### MAC5A physically interacts with ELP6

To explore the molecular function of MAC5A, we identified its interacting partners through a yeast two-hybrid (Y2H) screen of an Arabidopsis cDNA library. Among the candidates, ELP6 emerged as a high-confidence interactor. MAC5A harbors two structured domains, CCCH zinc-finger domain (ZnF_C3H1) and RRM (RNA recognizing motif). In contrast, ELP6 contains a single conserved ELP6 domain (Fig. 1a). To validate and map the interaction regions, we conducted Y2H assays using full-length and truncated variants of both proteins. The result showed that co-expression of the activation domain (AD)-fused MAC5A (AD-MAC5A) and DNA-binding domain (BD)-fused ELP6 (BD-ELP6) activated the expression of the reporter genes *HIS3*, *ADE2*, and *MEL1*, confirming their interaction in yeast (Fig. 1b). Domain dissection revealed that the fragment containing both the RRM domain and the C-terminal region of MAC5A is sufficient for binding ELP6 (Fig. 1b). Conversely, the ELP6 domain alone was insufficient for interaction with MAC5A, indicating that full-length ELP6 is required (Fig. 1b). We further validated this interaction in vitro using pull-down assays. MBP-tagged MAC5A specifically bound to GST-ELP6 immobilized on glutathione beads but not to GST control beads (Fig. 1c), confirming their directly physical interaction.

**Fig. 1.**
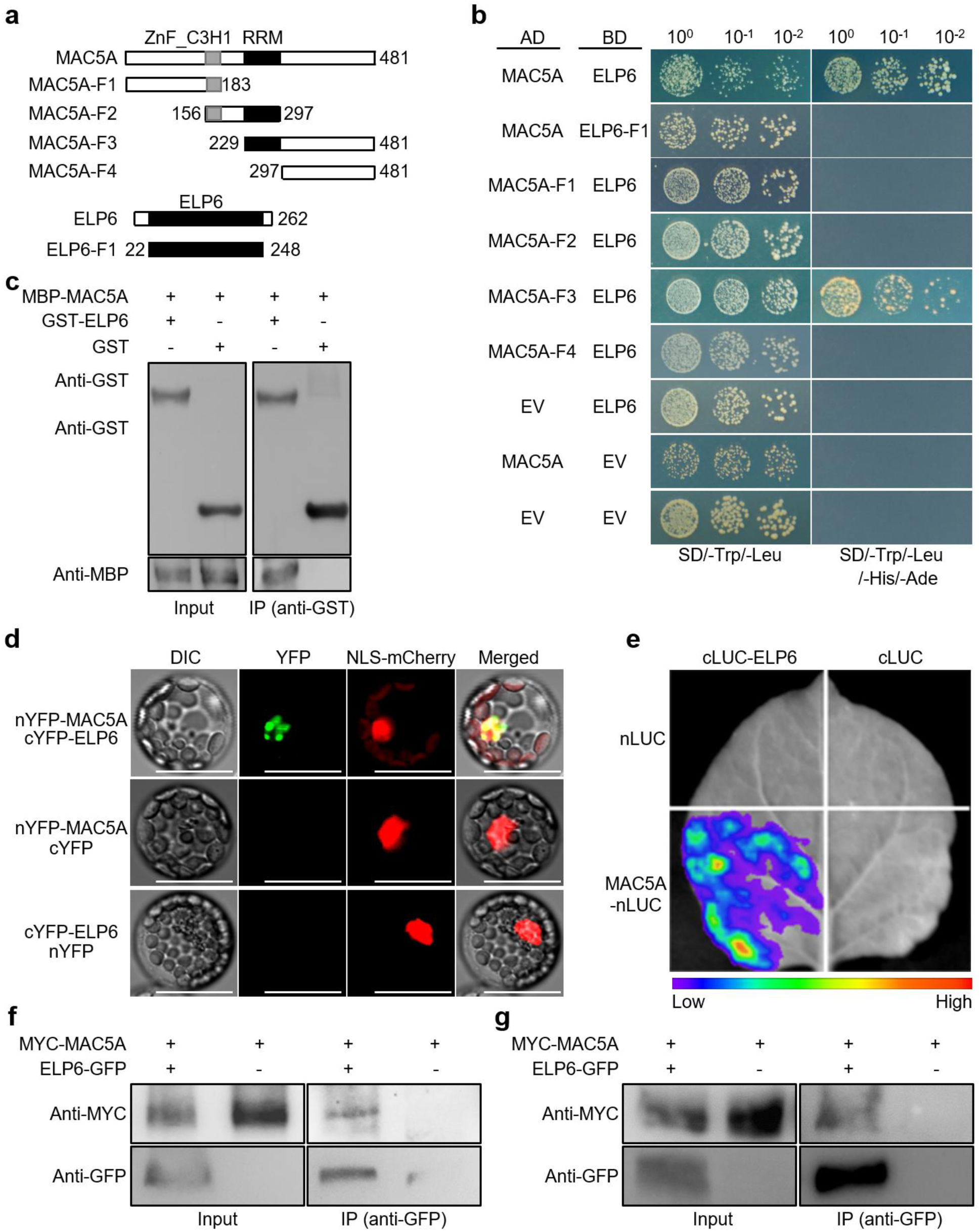
MAC5A physically interacts with ELP6. **a** Schematic diagrams of full-length and truncated MAC5A and ELP6 proteins. The conserved domains of MAC5A and ELP6, including ZnF_C3H1 (Zinc-Finger CCCH), RRM (RNA-recognition motif), and ELP6, are shown. **b** Y2H assay showing the interactions between MAC5A and ELP6. **c** Pull-down assay showing the physical interaction between MAC5A and ELP6 in vitro. **d** BiFC assay showing the interaction between MAC5A and ELP6 in Arabidopsis mesophyll protoplasts. The *35S*::*NLS-mCherry* plasmid was co-transformed to indicate the nucleus. Scale bars, 100 μm. **e** LCI assay showing the interaction between MAC5A and ELP6 in *N. benthamiana* leaves. **f** Co-IP assay showing the interaction between MAC5A and ELP6 in *N. benthamiana*. The plasmid combination of MYC-MAC5A and ELP6-GFP was transiently co-expressed in *N. benthamiana* leaves. IP was performed using anti-GFP antibody. After IP, MYC-MAC5A was detected by western blot using anti-MYC and anti-GFP antibodies. **g** Co-IP assay showing the interaction between MAC5A and ELP6 in stable transgenic Arabidopsis. The total proteins of 14d-old transgenic Arabidopsis plants harboring MYC-MAC5A and ELP6-GFP were extracted and Co-IP assay was performed. IP was performed using anti-GFP antibody. After IP, MYC-MAC5A was detected by western blot using anti-MYC and anti-GFP antibodies.

To confirm their interaction in plant cells, we examined subcellular localization of ELP6 by expressing ELP6-GFP in Arabidopsis mesophyll protoplast and the stable transgenic plants, respectively. ELP6-GFP fluorescence signal was detected within both nucleus and cytoplasm (Fig. S1). We subsequently employed bimolecular fluorescence complementation (BiFC) assays to assess the interaction between MAC5A and ELP6 in Arabidopsis protoplasts. Co-expression of nYFP-MAC5A and cYFP-ELP6 reconstituted YFP fluorescence exclusively in the nucleus, while control pairings yielded no signal (Fig. 1d). This finding was corroborated by Luciferase Complementation Imaging (LCI) in *Nicotiana benthamiana* leaves, where co-infiltration of MAC5A-nLUC and cLUC-ELP6 generated strong luminescence, absent in controls (Fig. 1e). Moreover, we transiently co-overexpressed the related construct pairs in *N. benthamiana* leaves and performed co-immunoprecipitation (Co-IP) assays. The results showed that MYC tagged-MAC5A protein (MYC-MAC5A) was immunoprecipitated by GFP-tagged ELP6 (ELP6-GFP) with GFP antibodies (Fig. 1f). Critically, Co-IP assay recapitulated MAC5A-ELP6 interaction in Arabidopsis plants harboring overexpressed ELP6-GFP and MYC-MAC5A (Fig. 1g). Taken together, these results provide a strong evidence that MAC5A physically interacts with ELP6.

### MAC5A functionally cooperates with ELP6 in plant development

Given the established roles of MAC5A and ELP6 in root and leaf development (23, 27), we investigated whether they functionally synergize to regulate plant growth. We first generated *mac5a elp6* double mutant by crossing *mac5a-2* and *elp6* alleles (Fig. S2). Both single mutants exhibited significantly shorter primary roots and reduced root apical meristem (RAM) cell numbers compared to wild-type (WT) plants (Fig. 2a-d). Strikingly, the *mac5a elp6* double mutant displayed synergistic defects, with further reductions in root length and RAM cell counts (Fig. 2a-d), suggesting their genetic interaction. To assess cell division activity, we introduced the *CycB1;1::GUS* and *CycB1;1::GFP* reporters (29, 30), markers of G2/M-phase progression, into the *mac5a-2* and *elp6* background, respectively. Both reporters showed markedly reduced expression in *mac5a elp6* roots compared to WT and single mutants (Fig. 2e), indicating that MAC5A and ELP6 cooperatively promote mitotic activity in the RAM. Additionally, the aerial tissue of *mac5a elp6* exhibited severe pleiotropic defects, including stunted vegetative growth, narrower rosette leaves, pin-like inflorescence architecture, disrupted apical-basal patterning of the gynoecium, and changed leaf venation patterns (Figs. 2f-i, S3). Additionally, since ELP6 is an accessory subunit of Elongator complex, we extended our analysis to its catalytic subunit, ELP1. The *mac5a elp1* double mutant was generated by crossing *mac5a-2* with *elp1-2* mutant, and phenotypic analyses indicated that *mac5a elp1* double mutant phenocopied the developmental defects of *mac5a elp6*, including synergistic reductions in root growth and meristem activity (Fig. S4), implying the involvement of both MAC5A and Elongator complex in apical meristem integrity.

**Fig. 2.**
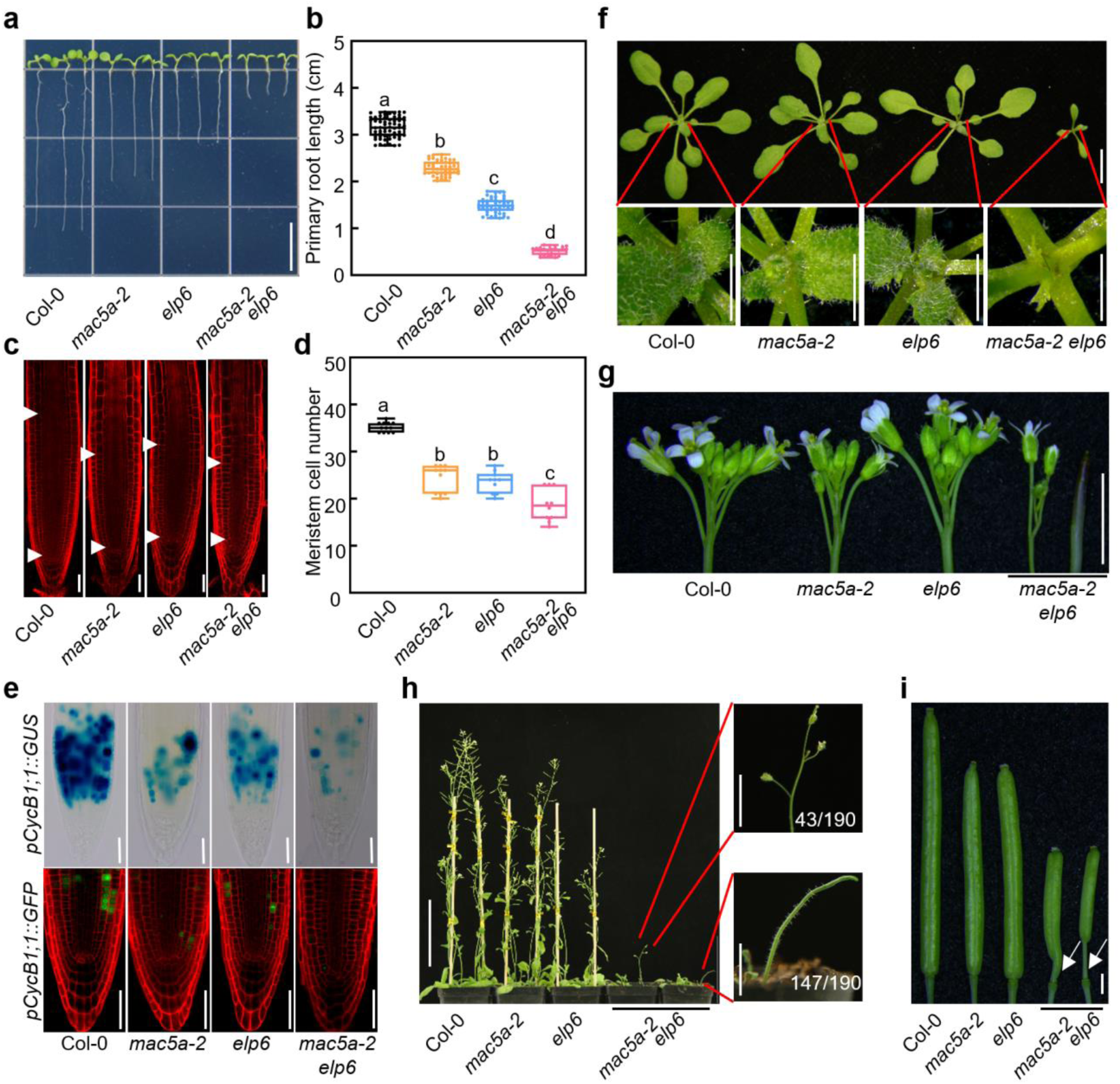
MAC5A acts with ELP6 to regulate plant development. **a** Representative images showing 7-day-old seedlings of Col-0, *mac5a-2*, *elp6*, and *mac5a-2 elp6*. Scale bar, 1 cm. **b** The statistical results of primary root length for Col-0, *mac5a-2*, *elp6*, and *mac5a-2 elp6* (n ≥ 25). Asterisks indicate significant differences determined by Student’s *t* test (** *P* < 0.01). **c** Representative images showing the RAM in 7-day-old seedlings of Col-0, *mac5a-2*, *elp6*, and *mac5a-2 elp6*. Propidium iodide (PI) staining was employed to visualize cell boundaries. White triangles indicate the meristem zone. Scale bars, 25 μm. **d** The statistical results of RAM cell numbers of Col-0, *mac5a-2*, *elp6*, and *mac5a-2 elp6* (n ≥ 10).. Different letters above the plots in (b, d) indicate significant differences (*P* < 0.05) according to one-way ANOVA with Tukey’s multiple comparisons test. **e** Analysis of *CYCB1;1* expression via *pCYCB1;1::GUS* staining (upper panels) and *pCYCB1;1::GFP* fluorescence (lower panels) in 5-day-old seedlings. Scale bars, 50 μm. **f** Representative images showing 3-week-old seedlings of Col-0, *mac5a-2*, *elp6*, and *mac5a-2 elp6*. The lower panels show the enlarged images of the shoot apical regions. Scale bars: 1 cm (upper panels); 1 mm (lower panels). **g** Inflorescence of 5-week-old plants in Col-0, *mac5a-2*, *elp6*, and *mac5a-2 elp6*. *mac5a-2 elp6* displays small or pin-like inflorescences. Scale bar, 1 mm. **h** Phenotype of 7-week-old plants of Col-0, *mac5a-2*, *elp6*, and *mac5a-2 elp6*. In the *mac5a-2 elp6* population, 23% (43/190) of plants produced inflorescences with flowers, while 77% (147/190) displayed pin-like inflorescences. Scale bars, 10 cm (left panel); 1 cm (right panel). **i** Siliques of Col-0, *mac5a-2*, *elp6-2*, and *mac5a-2 elp6-2*. Arrowheads highlight abnormal gynoecia. Scale bar, 1 mm.

### MAC5A and ELP6 are required for proper auxin distribution

The pronounced developmental defects in *mac5a elp6* prompted analysis of auxin distribution and signaling. We introduced the auxin response reporters *DR5::GUS* and *DR5::GFP* into the single and double mutant backgrounds of *mac5a* and *elp6*, respectively. Histochemical and fluorescence analyses revealed a substantial reduction in *DR5* activity in the root apices of the mutants, most notably in the *mac5a elp6* mutant, indicating a decrease in auxin response in these tissues (Fig. 3a and b). Given that leaves are a primary site of auxin biosynthesis, we next quantified auxin levels in the leaves of Col-0, *mac5a*, *elp6*, and *mac5a elp6* plants. Intriguingly, the auxin contents in leaf tissues were elevated in all tested mutants (Fig. 3c and d). Furthermore, the roots of these mutants, especially the *mac5a elp6* double mutant, displayed reduced sensitivity to L-kynurenine (Kyn), an inhibitor of auxin biosynthesis (Fig. 3e) (31), indicating the disrupted auxin homeostasis by the mutations of MAC5A and ELP6.

**Fig. 3.**
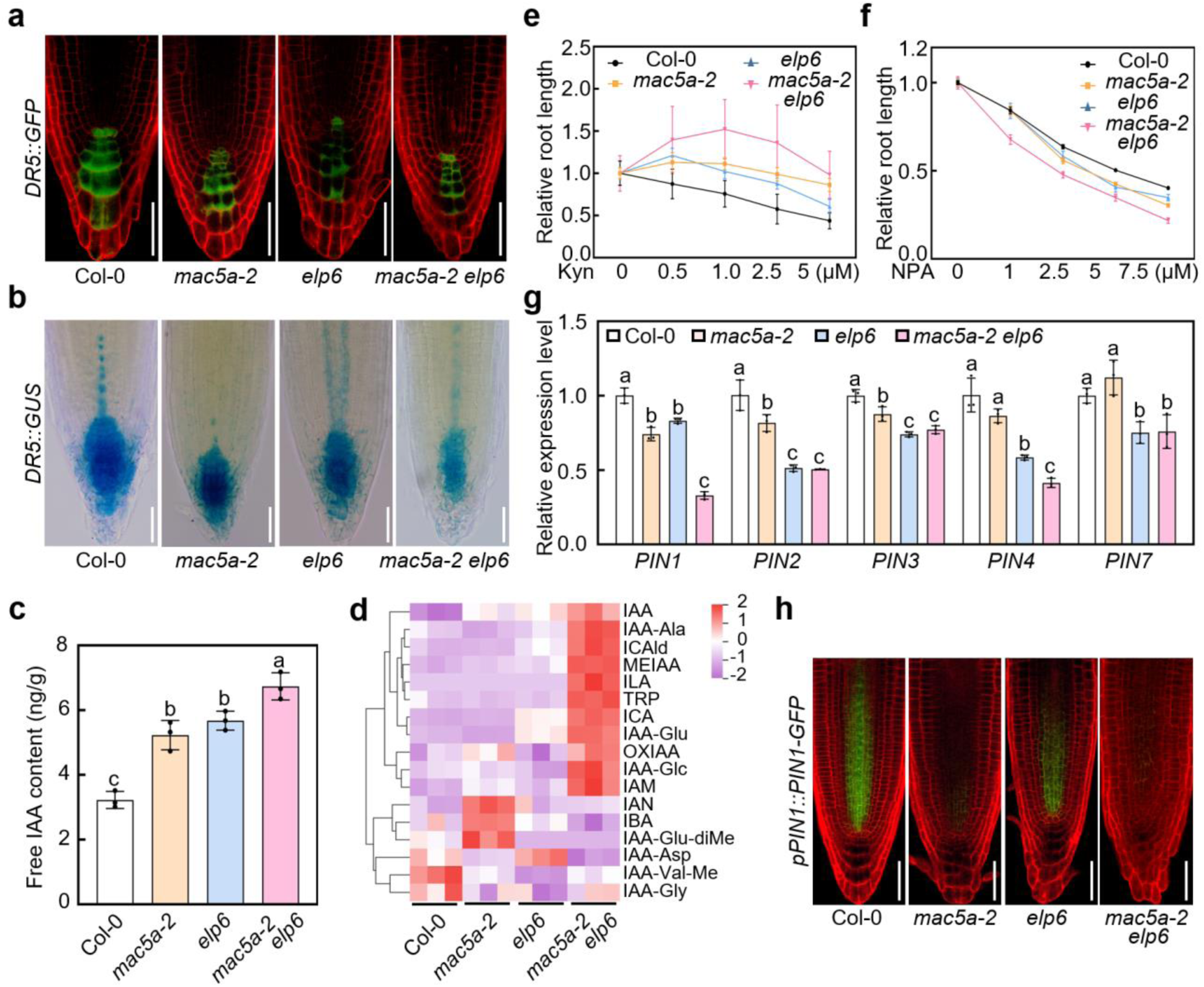
MAC5A and ELP6 affect auxin distribution. **a** Fluorescence signals of *DR5::GFP* in 5-day-old seedlings of Col-0, *mac5a-2*, *elp6*, and *mac5a-2 elp6.* PI staining was employed to visualize cell boundaries. Scale bars, 50 μm. **b** GUS staining of *DR5::GUS* in 5-day-old seedlings of Col-0, *mac5a-2*, *elp6*, and *mac5a-2 elp6*. Scale bars, 50 μm. **c** Free indole-3-acetic acid (IAA) content in Col-0, *mac5a-2*, *elp6*, and *mac5a-2 elp6*. Data are presented as mean ±SD (n = 3). Asterisks indicate significant differences using Student’s *t* test (** *P* < 0.01). **d** Heat map showing the content differences of auxin secondary metabolites in Col-0, *mac5a-2*, *elp6*, and *mac5a-2 elp6*. **e and f** The statistic results of relative primary root length of 7-day-old seedlings upon the treatments to Kyn (**e**) and NPA (**f**). Data are presented as mean ±SD (n ≥ 20). **g** RT-qPCR results showing the expression levels of *PIN* genes in 7-day-old seedlings of Col-0, *mac5a-2*, *elp6*, and *mac5a-2 elp6*. *ACTIN2* was used as the internal control. Data are presented as mean ± SD of three biological replicates. Different letters above the plots in (**c**, **g**) indicate significant differences (*P* < 0.05) according to one-way ANOVA with Tukey’s multiple comparison test. **h** Fluorescence signals of *pPIN1:PIN1-GFP* in 5-day-old seedlings of Col-0, *mac5a-2*, *elp6*, and *mac5a-2 elp6*. PI staining was emplolyed to visualize cell boundaries. Scale bars, 50 μm.

The impaired auxin distribution suggests that MAC5A and ELP6 may cooperately regulate auxin transport. To assess this possibility, we examined the sensitivity of the mutants to the polar auxin transport inhibitor N-1-naphthylphthalamic acid (NPA). Seedlings of all mutants, especially *mac5a elp6* double mutant, exhibited enhanced sensitivity to NPA treatment compared to WT (Fig. 3f), suggesting that MAC5A and ELP6 act synergistically to regulate auxin polar transportation. Polar auxin transport is predominantly facilitated by PIN efflux carriers. We therefore analyzed the expression of key *PIN* genes in root tips by RT-qPCR. Transcript levels of multiple *PIN* genes were significantly reduced in the mutants (Fig. 3g). Notably, the expression of *PIN1*, which encodes a major auxin efflux carrier critical for both shoot and root development, was most severely affected in the *mac5a elp6* mutant, indicating that MAC5A and ELP6 may act coordinately to regulate its transcription (Fig. 3g). Supporting this notion, PIN1-GFP accumulation is very low compared with WT and the single mutants (Fig. 3h). Collectively, these results suggest that MAC5A and ELP6 are required for proper auxin distribution by regulating *PIN* gene expression.

### MAC5A promotes transcription elongation of *PIN1* gene

Because MAC5A physically associates with Elongator (Fig. 1), a well-known regulator of transcriptional elongation and plant growth (21, 32), we hypothesized that MAC5A may play a role in the control of transcriptional elongation. To test this, we first assessed the sensitivity of plants to 6-Azauracil (6-AU), a chemical inhibitor of transcription elongation (33, 34). Wild-type plants exhibited a significant, dose-dependent repression of primary root growth on 6-AU (Fig. 4a and b). In contrast, both *mac5a-2* and *elp6* single mutants were less sensitive to this inhibition (Fig. 4a and b). Strikingly, the *mac5a elp6* double mutant was nearly insensitive to 6-AU (Fig. 4a and b), indicating that MAC5A and ELP6 synergistically regulate transcription elongation in planta. Since MAC5A is the accessory subunit of MAC complex, we further examined the sensitivity of mutants of the core subunits of MAC, including *mos4-2*, *cdc5-1*, *prl1-2*, and *mac3* to 6-AU. The analysis result demonstrated that like *mac5a-2*, all the tested mutants displayed decreased sensitivity compared with WT (Fig. S5). This result implies that the entire MAC complex may act in the control of transcriptional elongation.

**Fig. 4.**
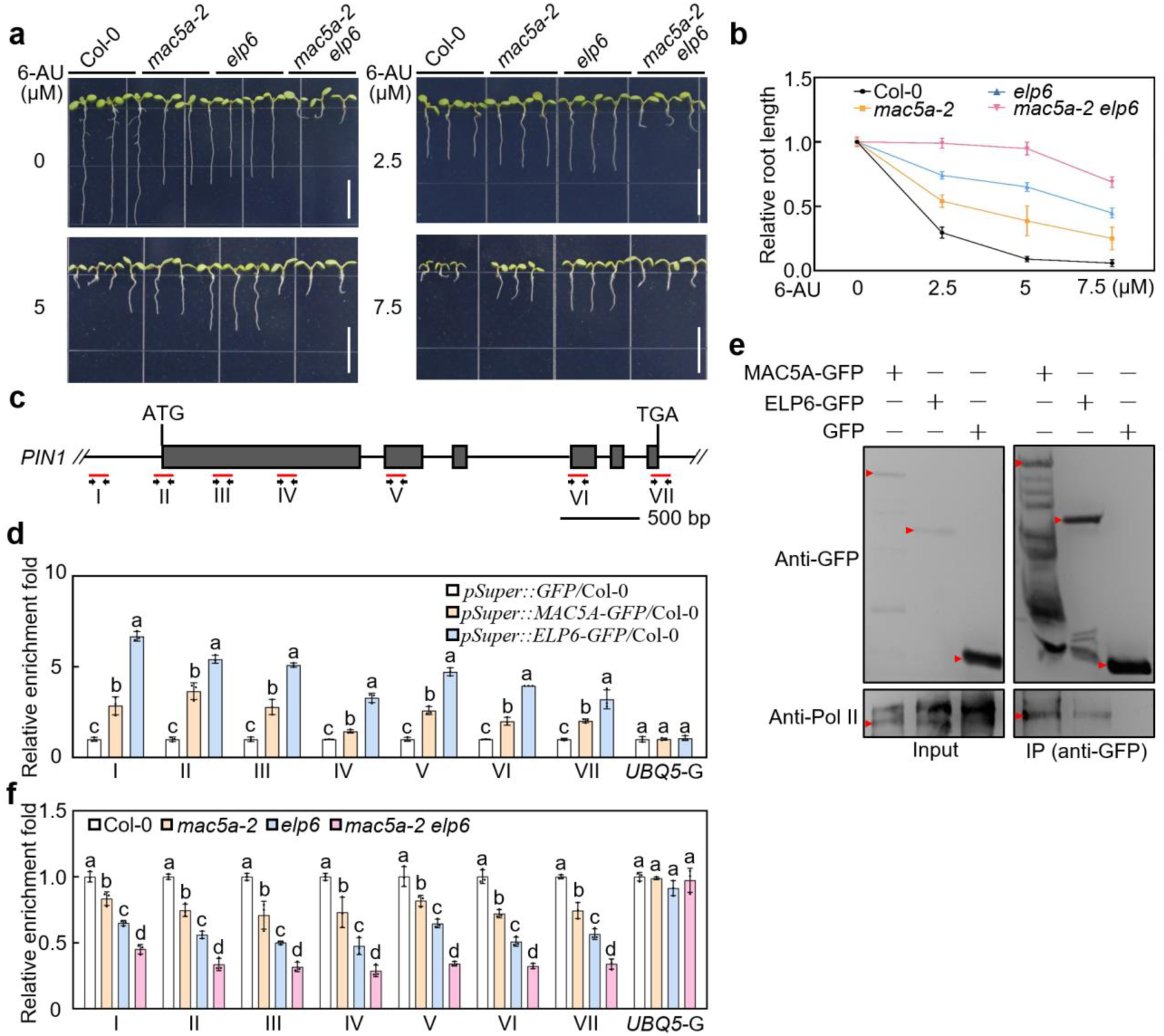
MAC5A and ELP6 synergistically regulate transcription elongation. **a** Representative images of 7-day-old seedlings of Col-0, *mac5a-2*, *elp6*, and *mac5a elp6* treated with 6-AU. Scale bars, 1 cm. **b** The statistical results of relative primary root length of 7-day-old seedlings upon 6-AU treatment. The values under normal condition were set as 1. Data are presented as mean ±SE (n ≥ 20). **c** Schematic diagram of *PIN1* gene structure. Regions used for ChIP-PCR were shown (I, II, III, IV, V, VI, and VII). Gray boxes indicate exons. **d** ChIP analysis results showing the enrichment of MAC5A and ELP6 on chromatin at the *PIN1* gene locus. ChIP assays were performed using 7-day-old seedlings. The values were calculated by normalizing immunoprecipitated DNA against input DNA. The values of *pSuper::GFP/*Col-0 were set as 1. *UBQ5* gene body (*UBQ5-G*) was employed as a negative control. Data are presented as mean ± SD. **e** Co-IP result showing the interactions between MAC5A or ELP6 with RNA polymerase II (Pol II). Total proteins were extracted from 14-day-old seedlings of *pSuper::MAC5A-GFP*, *pSuper::ELP6-GFP*, and *pSuper::GFP*, respectively. Immunoblotting analysis was performed with antibodies against GFP and CTD of Pol II. **f** ChIP analysis results showing the enrichment of phosphorylated RNA Pol II on chromatin at the *PIN1* gene locus in 7-day-old seedlings of Col-0, *mac5a-2*, *elp6*, and *mac5a elp6*. The values were calculated by normalizing immunoprecipitated DNA against input DNA. The values of Col-0 were set as 1. *UBQ5* gene body (*UBQ5-G*) was employed as a negative control. Data are presented as mean ±SD. Different letters above the plots in (d, f) indicate significant differences (*P* < 0.05) according to two-way ANOVA with Tukey’s multiple comparison test.

Since *PIN1* gene showed the reduced expression in *mac5a elp6* double mutant, we next asked whether MAC5A and ELP6 directly regulate its transcription. Chromatin immunoprecipitation (ChIP) assays revealed that both MAC5A and ELP6 are specifically enriched at the promoter and gene body regions of *PIN1* (Fig. 4c and d). Supporting this, Co-IP experiments confirmed that MAC5A and ELP6 interact with phosphorylated RNA polymerase II (Pol II) (Fig. 4e), the form essential for transcriptional elongation. To determine if the decreased *PIN1* transcript levels in the mutants resulted from a defect in elongation, we performed ChIP to monitor the occupancy of phosphorylated Pol II across the *PIN1* locus. The result demonstrated that Pol II association was dramatically reduced in both *mac5a-2* and *elp6* single mutants (Fig. 4f). Notably, this defect was further enhanced in the *mac5a elp6* double mutant (Fig. 4f). These results demonstrate that the MAC5A-ELP6 interaction is critical for the efficient recruitment or progression of elongating Pol II on the *PIN1* gene.

### The role of MAC5A partially dependents on functional ELP6

To define genetic hierarchy, we crossed *pSuper::ELP6-GFP/*Col-0 into *mac5a-2* and *pSuper::MAC5A-GFP/*Col-0 into *elp6* backgrounds. Overexpressing *ELP6* increased primary root length equally in both WT and *mac5a-2* (Fig. 5a and b), while *MAC5A* overexpression enhanced primary root length in WT but not *elp6* (Fig. 5c and d). Consistently, RT-qPCR results showed increased *PIN1* expression with *ELP6* overexpression in *mac5a-2*, but no change with *MAC5A* overexpression in *elp6* background (Fig. 5e and f). These data suggest that the role of MAC5A in root growth may partially dependent on functional ELP6 protein.

**Fig. 5.**
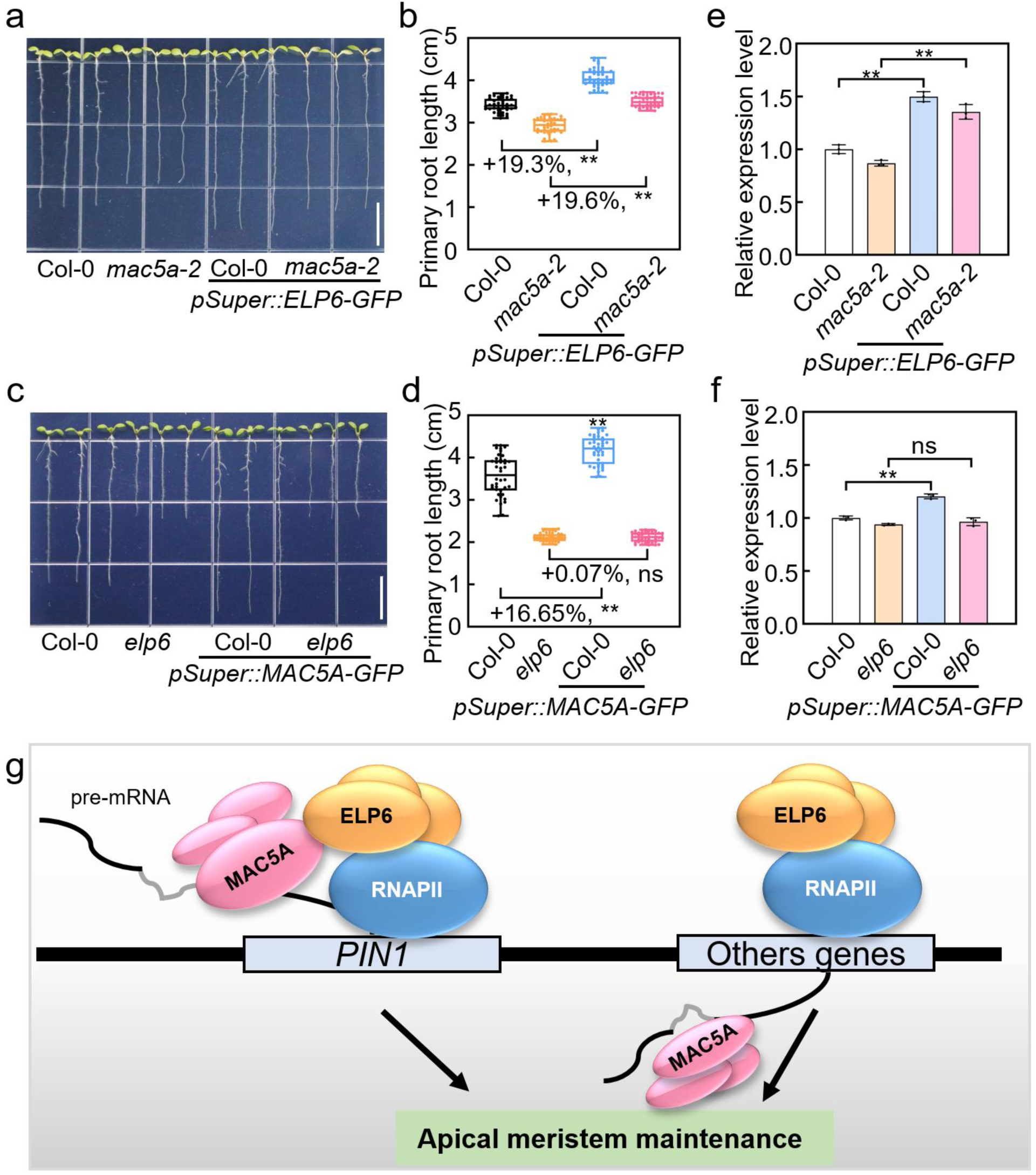
MAC5A acts upstream of ELP6 to regulate *PIN1* transcription. **a and b** Representative images (**a**) and statistical results (**b**) of primary root length of 7-day-old seedlings of Col-0, *mac5a-2*, *pSuper::ELP6-GFP/*Col-0, and *pSuper::ELP6-GFP/mac5a-2.* **c and d** Representative images (**c**) and statistical results (**d**) of primary root length of 7-day-old seedlings of Col-0, *elp6*, *pSuper::MAC5A-GFP/*Col-0, *pSuper::MAC5A-GFP/elp6*. Scale bars in **a** and **c**, 1 cm. Data are presented as mean ± SD (n ≥ 40). **e and f** RT-qPCR results showing *PIN1* expression in 7-day-old seedlings. *ACTIN2* was used as the internal control. Data are presented as mean ±SD of three biological replicates. Asterisks in **b**, **d**, **e**, and **f** indicate significant differences using Student’s *t* test (** *P* < 0.01, ns: no significance). **g** A proposed model illustrates the synergistic role of MAC5A and ELP6 in maintaining meristem integrity. During the transcription elongation of PIN1, MAC5A directly interacts with ELP6, and both associate with RNAPII CTD to modulate the elongation efficiency of PIN1 transcription viaELP6. In parallel, MAC5A and ELP6 may execute independent functions in the control of gene expression. Black or gray line indicates exon or intron, respectively.

## Discussion

The coupling of mRNA splicing to transcription elongation is emerging as a central principle of eukaryotic gene regulation (35, 36), yet the molecular basis of this coordination in plants remain poorly defined. In this study, we provide direct functional and physical evidence that the splicing-associated factor MAC5A connects MAC complex to the transcriptional Elongator complex, enabling precise control of *PIN1* expression and meristem maintenance (Fig. 5g).

Our finding establish MAC5A as a crucial adaptor that bridges RNA-processing machinery with the transcriptional apparatus. Through its interaction with the Elongator subunit ELP6, MAC5A enhances RNA polymerase II occupancy and elongation at the *PIN1* locus, a central regulator of auxin flux and developmental patterning (2, 6, 8). Genetic analyses further demonstrate that MAC5A and Elongator synergistically support apical meristem activity, and reduced sensitivity to the elongation inhibitor 6-AU in multiple MAC mutants suggests that the entire MAC complex contributes to the elongation control.

The molecular architecture of MAC5A offers insights into its role. Its conserved ZnF_C3H1 and RRM domains likely confer RNA-binding capability, whereas the C-terminal region mediates ELP6 association. This organization positions MAC5A as an integrative hub capable of coupling nascent transcripts with elongating RNAPII. Notably, its human homolog RBM22 interacts with transcription elongation factor SPT5 to modulate RNAPII dynamics (37), pointing to an evolutionarily conserved strategy by which RNA-binding proteins govern transcriptional elongation across eukaryotes. Supporting this idea, the RNA-binding protein MINIYO (IYO) in Arabidopsis and its mammalian counterpart RPAP1 also interface with Elongator to drive stem cell differentiation (38, 39). Unlike MAC5A, IYO promotes differentiation through a distinct cytosol-to-nucleus translocation mechanism. However, the specific roles of the RNA-binding domain and its associated RNAs in modulating elongation remain to be elucidated.

The MAC5A-ELP6 interaction may also influence Elongator stability and recruitment. As a structural core subunit, ELP6 could enable MAC5A to help assemble or guide Elongator-RNAPII complexes to specific genomic targets such as *PIN1*. In addition, since Elongator promotes histone H3 acetylation (40), MAC5A-Elongator cooperation may help maintain an open chromatin state conductive to efficient elongation. On the RNA-processing side, MAC plays an essential role in pre-mRNA splicing to ensure sufficient level of functional transcripts. It is conceivable that the efficient intron removal could increase the elongation rate (16), likely via a MAC5A-mediated manner. This hypothesis is supported by our genetic evidence that MAC5A acts upstream of ELP6 to activate *PIN1* expression. Moreover, while our data firmly establish a nuclear role for MAC-Elongator interaction in transcriptional elongation, Elongator’s pleiotropic functions must be considered. In the cytoplasmic, Elongator’s role in tRNA modification likely affects translation and auxin response. These nuclear and cytoplasmic functions may operate independently yet converge on shared developmental outcomes.

In summary, this study identifies MAC5A as a molecular integrator that links co-transcriptional RNA processing with transcriptional elongation via the Elongator complex. This MAC5A-Elongator module orchestrates *PIN1* transcription and auxin transport to preserve meristem activity and plant architecture. Our work establishes a conceptual framework for the coupling of RNA processing to transcriptional elongation in plants and suggests an evolutionarily conserved principles of gene expression regulation across eukaryotes.

## MATERIALS AND METHODS

### Plant Materials and Growth Conditions

All *Arabidopsis thaliana* lines used in this study are in the Columbia-0 (Col-0) ecotype background. The *mac5a-2* (SALK_142085) T-DNA insertion mutant was obtained from the Arabidopsis Biological Resource Center (ABRC) and has been previously described (28). The *elp6* mutant was also as previously reported (23). The *mac5a-2 elp6* double mutant was generated by crossing *mac5a-2* with *elp6*. *pSuper::ELP6-GFP/mac5a-2* transgenic plant was produced by crossing *pSuper::ELP6-GFP/Col-0* with *mac5a-2*, and *pSuper::MAC5A-GFP/elp6* transgenic plant was produced by crossing *pSuper::MAC5A-GFP/Col-0* with *elp6*. Homozygous line was isolated from the F2 progeny by genotyping. The complementation line *35S::MYC-MAC5A/mac5a-2 (MAC5Acom-1)* was as reported (28).

Seeds were surface-sterilized with 10% (v/v) sodium hypochlorite for 6 minutes, followed by a brief rinse with 75% (v/v) ethanol and four rinses with sterile distilled water. Seeds were sown on half-strength Murashige and Skoog (1/2 MS) medium (Murashige et al., 1962), pH5.7, supplemented with 1% (w/v) sucrose and solidified with 0.8% (w/v) agar. Plates were stratified at 4°C in darkness for 2 days and then transferred to a controlled-environment growth chamber (Greenfuture Envirotech). Plants were grown under long-day conditions (16 h light/8 h dark cycle) at 22°C, with a light intensity of 120 µmol m^-2^ s^-1^ and 65% relative humidity. Seven-day-old seedlings were transferred to soil and maintained under the same conditions.

### Plasmid Constructions and Plant Transformation

To generate *pSuper::MAC5A-GFP* and *pSuper::ELP6-GFP* constructs, the full-length coding sequence (CDS) of *MAC5A* and *ELP6* were amplified from Arabidopsis Col-0 using gene-specific primers (Supplemental Table S1). The PCR products were cloned into the *pCambia1300* vector, respectively. For plant transformation, recombinant plasmids were introduced into *Agrobacterium tumefaciens* strain *GV3101* by electroporation. Arabidopsis plants (Col-0, *mac5a-2*, or *elp6*) were transformed using the floral dip method (41). Transgenic lines were selected on 1/2 MS medium containing 50 mg L^-1^ hygromycin.

### Y2H Assay

Protein-protein interactions were conducted using the MATCHMAKER GAL4 Two-Hybrid System (TaKaRa, Cat. 630487). For the initial screen, the CDS of MAC5A was cloned into the *pGBKT7* bait vector. To confirm interactions and map domains, full-length and truncated versions of MAC5A and ELP6 were cloned into *pGADT7* and *pGBKT7* vectors, respectively. Construct pairs were co-transformed into the yeast strain *Y2Hgold* (Clontech) using the lithium acetate method. Transformants were selected on synthetic dropout (SD) medium lacking leucine and tryptophan (SD/-Trp /-Leu). Protein interactions were assessed by growth on high-stringency SD medium lacking leucine, tryptophan, histidine, and adenine (SD/-Trp/-Leu/-His/-Ade).

### BiFC Assay

The CDS of *MAC5A* was cloned into *pVYNE (R)* vector to generate *35S::nYFP-MAC5A*. The CDS of *ELP6* was cloned into *pVYCE (R)* vector to generate *35S::cYFP-ELP6* (Waadt et al., 2008). These constructs, along with a nucleus marker plasmid (*35S::NLS-mCherry*), were co-transfected into Arabidopsis mesophyll protoplasts via polyethylene glycol (PEG)-mediated transformation (42). The combinations of *35S::nYFP-MAC5A* with empty *35S::pVYCE (R)*, *35S::cYFP-ELP6* with empty *35S::pVYNE (R)*, and *35S::pVYCE (R)* with *35S::pVYNE (R)* were employed as negative controls. After incubation in the dark at 22°C for 12 h, YFP and mCherry fluorescence was detected using a confocal laser scanning microscope (Olympus, FluoView FV1000). YFP was excited at 488 nm with emission collected at 500–540 nm; mCherry was excited at 561 nm with emission collected at 570–620 nm.

### LCI Assays

The CDS of *MAC5A* was cloned into *pCambia1300-nLUC* vector to generate *35S::MAC5A-nLUC*. The CDS of *ELP6* was cloned into *pCambia1300-cLUC* vector to generate *35S::cLUC*-*ELP6-* (Waadt et al., 2008). Constructs were transformed into *A.tumefaciens* strain *GV3101*. Bacterial cultures harboring the respective constructs were mixed 1:1 (v/v) and co-infiltrated into the leaves of *N. benthamina* plants. The combinations of *35S::MAC5A-nLUC* with empty *35S::cLUC*, *35S::cLUC-ELP6* with empty *35S::nLUC*, and empty *35S::nLUC* with *35S::cLUC* were employed as negative controls. Infiltrated plants were kept in darkness for 24 h and then under a 16 h light/8 h dark cycle for 24 h. Luciferase activity was detected by spraying leaves with 1 mM D-luciferin and imaging with a NIGHTSHADE LB985 in vivo imaging system (Berthold Technologies).

### Co-IP Assays

The CDS of *MAC5A* was cloned into a modified *PBI221* vector to generate *35S::MYC-MAC5A*. This construct was co-infiltrated with *pSuper::ELP6-GFP* into *N. benthamiana* leaves via *Agrobacterium*-mediated transformation. After 40 h, leaf tissue was harvested and total protein was extracted with extraction buffer (50 mM Tris-HCl, pH 7.6, 150 mM NaCl, 5 mM MgCl_2_, 0.5 mM dithiothreitol (DTT), 10% glycerol, 0.1% Nonidet P-40, 1 mM phenylmethanesulfonyl fluoride (PMSF), and 1× protease inhibitor cocktail). The supernatant was incubated with anti-MYC agarose beads (Myc-Trap® Agarose, Proteintech) for 2 h at 4°C with gentle rotation. After incubation, beads were washed five times with washing buffer (50 mM Tris-HCl, pH 7.5, 150 mM NaCl, 0.1% Triton X-100, 20% glycerol, 1 mM EDTA, 1 mM PMSF, and 1× protease inhibitor cocktail). Bound proteins were eluted by boiling in 2× SDS loading buffer for 10 min, separated by SDS-PAGE, and immunoblotted using anti-GFP (TransGen, HT801) and anti-MYC (TransGen, HT101) antibodies, respectively. For the protein interaction between phosphorylated Pol II and MAC5A or ELP6, total proteins were extracted from 14-day-old seedlings of *pSuper::MAC5A-GFP*, *pSuper::ELP6-GFP*, and *pSuper::GFP*, respectively. Immunoblotting analysis was performed with antibodies against GFP (TransGen, HT801) and CTD of Pol II (Abcam, ab5131).

### GST Pull-down Assay

The CDS of *MAC5A* was cloned into *pETMALc-H* for expression as an MBP fusion protein. The CDS of *ELP6* was cloned into *pGEX-4T-1* for expression as a GST fusion protein. Recombinant plasmids were transformed into *Escherichia coli* BL21 (DE3) cells. Protein expression was induced with 0.5 mM isopropyl β-D-1-thiogalactopyranoside (IPTG) at 16°C overnight. MBP-MAC5A fusion protein was purified using amylose resin (BioLabs, E8021V). GST-ELP6 and GST alone were purified using Glutathione Sepharose 4B (Cytiva, 17527901). For pull-down assays, purified MBP-MAC5A was incubated with GST-ELP6 or GST bound to glutathione beads for 1 h at 4°C. Beads were washed five times with PBS buffer, and bound proteins were eluted with elution buffer (50 mM Tris-HCl, pH 7.4, 150 mM NaCl, 0.5% Triton X-100, 5% glycerin and 100 mM reduced glutathione). Elutes were analyzed by immunoblotting using anti-MBP (TransGen, HT701) and anti-GST (CWBIO, CW0084M) antibodies, respectively.

### RNA Extraction and qRT-PCR Analysis

Total RNA was isolated from seven-day-old seedling using the TransZol Up RNA Kit (TransGen, ET111). First-strand cDNA was synthesized from 1 µg RNA using the TransScript One-Step gDNA Removal and cDNA Synthesis SuperMix (TransGen, AT311). Quantitative RT-PCR (qRT-PCR) was performed with TransStart Top Green qPCR SuperMix (TransGen, AQ131) on a QuantStudio One Real-Time PCR System (Thermo Fisher Scientific, USA). *ACTIN2* was used as an internal reference gene. Relative expression levels were calculated using the 2^-ΔΔCT^ method (43). All primer sequences are listed in Supplemental Table S1.

### Propidium iodide (PI) Staining and Confocal Microscopy

Five-day-old seedlings were immersed in 10 μg mL^−1^ propidium iodide (PI) solution for 1 min and rinsed with deionized water. PI staining (excitation 559 nm, emission 600–650 nm) and GFP/RFP fluorescence (GFP: excitation 488 nm, emission 500–540 nm; RFP: excitation 558 nm, emission 580–630 nm) were visualized using a confocal laser scanning microscope (Olympus Fluo View FV1000).

### Histochemical GUS Staining

Five-day-old seedlings harboring the GUS reporter gene were incubated in GUS staining buffer (50 mM Na_3_PO_4_, pH 7.0, 0.1% TritonX-100, 0.4 mM K_4_[Fe(CN)_6_]·3H_2_O, 0.4 mM K_3_[Fe(CN)_6_], 0.5 mg/mL X-glucuronide) at 37°C for approximately 6 h. The stained tissues were cleared by sequential washes with 70% ethanol. Cleared tissues were photographed under a stereomicroscope (Leica, MZ FLIII).

### ChIP Assay

ChIP assays were performed as described (44, 45) using seven-day-old seedlings. Approximately 1 g of tissues were cross-linked with 1% (v/v) formaldehyde under vacuum. Chromatin was isolated, sonicated to an average fragment size of 100–500 bp, and immunoprecipitated overnight at 4°C with anti-GFP antibody (Abcam, ab290). After reverse cross-linking and DNA purification, enriched DNA fragments were quantified by qPCR using gene-specific primers (Supplemental Table S1).

### Statistical Analysis

All experiments were performed with at least three independent biological replicates. Data are presented as mean ± standard deviation (SD). Statistical significance was determined using Student’s *t* test or one-way ANOVA in GraphPad Prism 9 software.

## SUPPLEMENTAL DATA

**Supplemental Figure 1. ELP6 localizes in nucleus and cytoplasm.**

**Supplemental Figure 2. Genotyping of *elp6* mutant.**

**Supplemental Figure 3. MAC5A and ELP6 regulate leaf development.**

**Supplemental Figure 4. MAC5A acts with ELP1 to regulate plant development.**

**Supplemental Figure 5. MAC complex regulates transcript elongation.**

**Supplemental Table S1. Primers used in this study.**

## Supporting information

supplemental data set

## ACKNOWLEDGMENTS

We thank Professor Zhaojun Ding (Shandong University) for providing *elp6* seeds. We also acknowledge Professor Yunhai Li (Institute of Genetics and Developmental Biology, Chinese Academy of Sciences) for providing the seeds of *CYCB1;1::GFP*, *CYCB1;1::GUS, DR5::GFP*, *DR5::GUS*, and *PIN1::PIN1-GFP*. This study was financially supported by the Key R&D Program of Shandong Province, China (2026CXPT044), National Natural Science Foundation of China (No. 32070621, No. 32470317, and No. 32670764), and Double First-Class Discipline Construction Fund of Shandong Agricultural University.

## AUTHOR CONTRIBUTIONS

Shengjun Li designed the research. Meng Ye, Xudong Li, Yufeng Zhou, Xiaojuan Huang, Huilin Liu, and Shuqing Liu performed the experiments and analyzed the data. Ruibo Hu and Aixia Li analyzed the data and discussed the results. Shengjun Li and Meng Ye wrote the article with contributions from all authors.

## Competing interests

The authors declare no competing interesting.

## Notes

### Competing Interest Statement

The authors have declared no competing interest.

