## supplemental data set for "The MOS4-associated complex subunit MAC5A maintains meristem development by regulating transcription elongation in Arabidopsis"

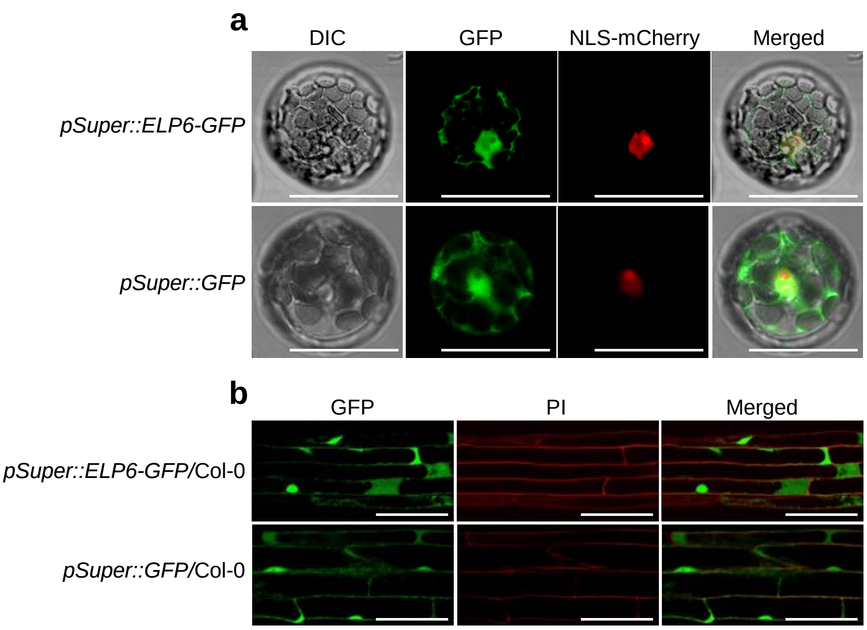


**Supplementary Fig. 1 ELP6 localizes in nucleus and cytoplasm.**

**a,** Subcellular localization of ELP6 in Arabidopsis mesophyll protoplasts. The plasmid combinations of *pSuper::ELP6-GFP* or *pSuper::GFP* with *35S*::*NLS-mCherry* were introduced into protoplasts using PEG-mediated transformation. NLS-mCherry protein was employed to indicate the nucleus. Scale bars, 100 μm. **b,** Subcellular localization of ELP6-GFP or GFP in stable transgenic Arabidopsis plants. Roots of 5-day-old seedlings were used to observe the ELP6-GFP or GFP fluorescence signals. PI staining was emplolyed to visualize cell boundaries. Scale bars, 50 μm.

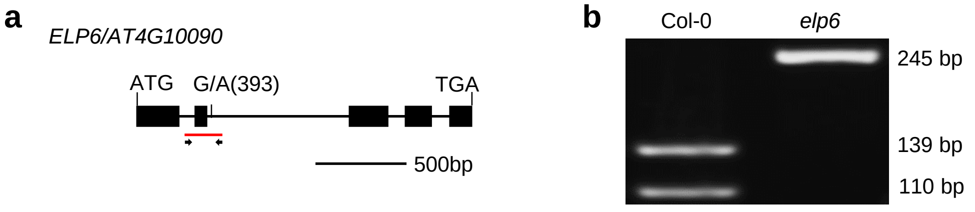


**Supplementary Fig. 2 Genotyping of *elp6* mutant.**

**a,** Schedemic diagram of *ELP6* gene. *elp6* mutant harbors a site mutation from G to A. The positions of primers used for dCAPS PCR were shown. **b,** Genotyping of *elp6* mutant using dCAPS PCR. PCR products from wild-type (Col-0), but not the *elp6* mutant, were diagested by *Bsm*AI enzyme.

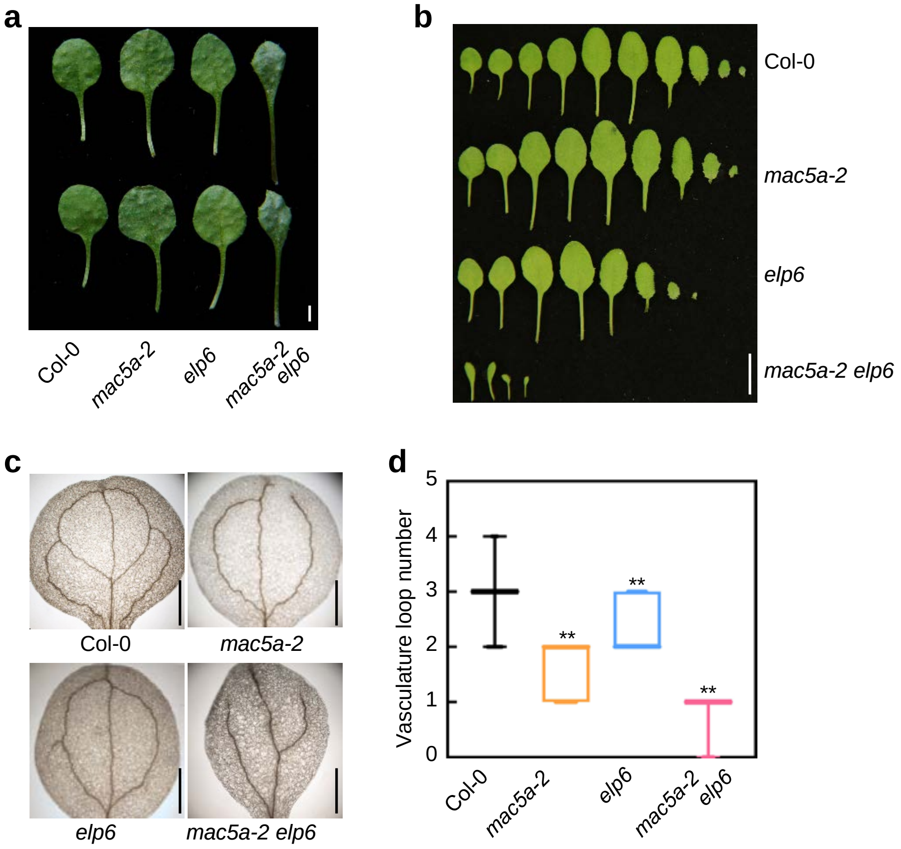


**Supplementary Fig. 3 MAC5A and ELP6 regulate leaf development.**

**a,** The first-pair of leaves from 3-week-old plants of Col-0, *mac5a-2*, *elp6,* and *mac5a-2 elp6*. Scale bars, 2 mm. **b,** Leaves from 3-week-old plants of Col-0, *mac5a-2*, *elp6,* and *mac5a-2 elp6*. Scale bars, 1 cm. **c,** The representative images showing cotyledons of Col-0, *mac5a-2*, *elp6,* and *mac5a-2 elp6.* Cotyledons were clarified with ethanol and subsequently mounted on slides for microscopic analysis. Scale bars, 1 mm. **d,** The statistical results of vasculature loop number of the cotyledons in Col-0, *mac5a-2*, *elp6,* and *mac5a-2 elp6* (n ≥ 30). Asterisks indicate significant differences using Student's *t* test (** $P < 0.01$).

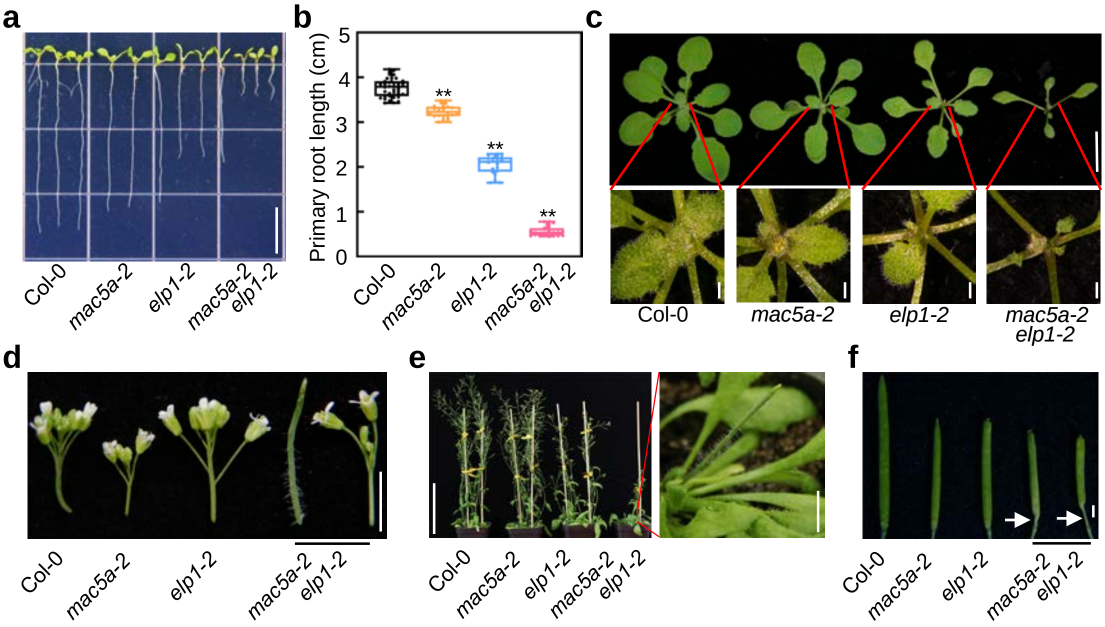


**Supplementary Fig. 4 MAC5A acts with ELP1 to regulate plant development.**

**a,** The representative images showing 7-day-old seedlings of Col-0, *mac5a-2*, *elp1-2,* and *mac5a-2 elp1-2*. Scale bar, 1 cm. **b,** The statistical results of primary root length of Col-0, *mac5a-2*, *elp1-2,* and *mac5a-2 elp1-2* (n ≥ 20). Asterisks indicate significant differences using Student's *t* test (** $P < 0.01$). **c,** The representative images showing 3-week-old seedlings of Col-0, *mac5a-2*, *elp1-2,* and *mac5a-2 elp1-2*. The bottom panels show the enlarged images of the shoot apical regions. Scale bars, 1 cm for upper; 1 mm for bottom. **d,** Inflorescences of 5-week-old plants in Col-0, *mac5a-2*, *elp1-2,* and *mac5a-2 elp1-2*. *mac5a-2 elp1-2* displays the smaller or pin-like inflorescences. Scale bar, 0.5 cm. **e,** 7-week-old plants of Col-0, *mac5a-2*, *elp1-2,* and *mac5a-2 elp1-2*. Scale bars, 10 cm for left; 1 cm for right. **f,** Siliques of Col-0, *mac5a-2*, *elp1-2,* and *mac5a-2 elp1-2*. Arrowheads indicate the abnormal gynoecium. Scale bar, 1 mm.

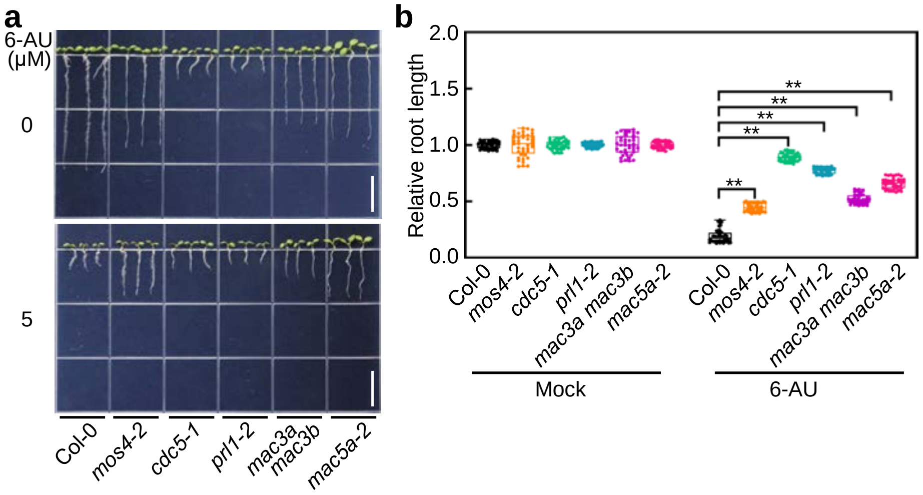


**Fig. S5 | MAC complex regulates transcript elongation.**

**a** The representative images of 7-day-old seedlings of Col-0, *mos4-2*, *cdc5-1*, *prl1-2*, *mac3a mac3b,* and *mac5a-2* in the response to 6-AU treatment. Scale bars, 1 cm. **b** The statistical results of relative primary root length of 7-day-old seedlings upon 6-AU treatment. Data are presented as mean ± SE ( n ≥ 40). The values under mock condition were set as 1. Asterisks indicate significant differences using Student's *t* test (** *P* < 0.01).
